# Repeated shifts in sociality are associated with fine-tuning of highly conserved and lineage-specific enhancers in a socially flexible bee

**DOI:** 10.1101/2024.08.15.608154

**Authors:** Beryl M. Jones, Andrew E. Webb, Scott M. Geib, Sheina Sim, Rena M. Schweizer, Michael G. Branstetter, Jay D. Evans, Sarah D. Kocher

## Abstract

Comparative genomic studies of social insects suggest that changes in gene regulation are associated with evolutionary transitions in social behavior, but the activity of predicted regulatory regions has not been tested empirically. We used STARR-seq, a high-throughput enhancer discovery tool, to identify and measure the activity of enhancers in the socially variable sweat bee, *Lasioglossum albipes*. We identified over 36,000 enhancers in the *L. albipes* genome from three social and three solitary populations. Many enhancers were identified in only a subset of *L. albipes* populations, revealing rapid divergence in regulatory regions within this species. Population-specific enhancers were often proximal to the same genes across populations, suggesting compensatory gains and losses of regulatory regions may preserve gene activity. We also identified 1182 enhancers with significant differences in activity between social and solitary populations, some of which are conserved regulatory regions across species of bees. These results indicate that social trait variation in *L. albipes* is driven both by the fine-tuning of ancient enhancers as well as lineage-specific regulatory changes. Combining enhancer activity with population genetic data revealed variants associated with differences in enhancer activity and identified a subset of differential enhancers with signatures of selection associated with social behavior. Together, these results provide the first empirical map of enhancers in a socially flexible bee and highlight links between *cis*-regulatory variation and the evolution of social behavior.

## Introduction

Social insects provide a textbook example of how changes in gene regulation can generate diverse phenotypes. Within their eusocial societies, overlapping generations of reproductive queens and non-reproductive workers cooperate as a group to reproduce collectively. Remarkable phenotypic plasticity is encoded within the social insect genomes because nearly any fertilized egg has the potential to develop into either a queen or a worker. Within the colony, queens specialize on reproductive tasks such as egg laying, while workers often specialize in caring for young, foraging, or defense (Michener 1974). Numerous studies have identified transcriptional (Evans and Wheeler 1999; Pereboom et al. 2005; Feldmeyer et al. 2014; Jones et al. 2017) and epigenetic differences (Herb et al. 2012; Weiner et al. 2013; Yan et al. 2014; Simola et al. 2015; Wojciechowski et al. 2018) between social insect castes. As a result, gene regulation is thought to play an essential role in the evolution of eusociality.

Among insects, eusociality has evolved from solitary ancestors at least 18 times (Bourke 2011), enabling studies of the convergent mechanisms of social evolution. Comparative genomic studies in social insects found support for a role of gene regulatory evolution in social origins, with predicted transcription factor (TF) binding presence expanded in the genomes of social compared with solitary bees (Kapheim et al. 2015; Jones et al. 2023) and high divergence rates of TF binding sites among ant species (Simola et al. 2013). In addition, changes in the evolutionary rates of predicted regulatory regions are associated with both the origins and maintenance of sociality (Rubin et al. 2019; Jones et al. 2023). These studies suggest that at the species level, social traits evolve alongside changes in gene regulation. However, we still lack understanding of how gene regulatory variation mediates intraspecific differences in social behavior, and how this variation may be selected during evolutionary transitions in sociality.

Species harboring natural variation in social behavior provide an excellent opportunity to study the role of gene regulation in mediating the evolution of social traits. *Lasioglossum albipes* is a socially flexible sweat bee species (Hymenoptera: Halictidae) where multiple populations of this bee have convergently reverted from eusociality to a solitary life history (Plateaux-Quenu 1993; Plateaux-Quénu et al. 2000; Kocher et al. 2018). Previous work demonstrates a genetic component underlies this social variability, and genetic variants associated with social status are often located in non-coding regions of the genome (Kocher et al. 2018), making *L. albipes* an ideal system to study the contribution of regulatory variation to social evolution.

One mechanism by which gene regulatory changes can evolve is via modifications to transcriptional enhancers. Enhancers are regulatory sequences that can modulate expression levels of associated genes, and they can be located upstream, downstream, or even within the introns of the genes they regulate (Maston et al. 2006; Levine 2010; Yáñez-Cuna et al. 2013). Enhancers are often tissue– or condition-specific (Yáñez-Cuna et al. 2012; Arnold et al. 2013) and can act over long distances (Lettice 2003). These regulatory elements serve as binding sites for TFs (Istrail and Davidson 2005; Levine 2010; Yáñez-Cuna et al. 2013) and mutations in enhancer sequences can alter their binding affinities (Gompel et al. 2005; Bradley et al. 2010; Reddy et al. 2012; Lim et al. 2024).

Enhancers are rapidly evolving, typically changing at faster rates than the genes they regulate or the TFs they interact with (Thurman et al. 2012). As a result, it is common for enhancers to exhibit rapid turnover across species (Arnold et al. 2014; Villar et al. 2015) and even across different populations within a species (Lewis and Reed 2019). Despite their relatively rapid sequence evolution, enhancer functions can be highly conserved across hundreds of millions of years (Wong et al. 2020). Over evolutionary time, enhancers can also be repurposed to regulate different genes, or new enhancers can arise from previously non-regulatory sequences (Long et al. 2016).

The role of enhancers in evolution has perhaps been best studied in the context of novel morphological traits (Carroll 2000; Hoekstra and Coyne 2007; Prud’homme et al. 2007; Carroll 2008). For example, the loss of trichomes in *Drosophila sechellia* larvae evolved via loss of enhancer elements of the TF *shavenbaby* (*svb*) (Frankel et al. 2011), and wing pattern variation in *Heliconius* butterflies involves enhancer modifications that rewire the gene regulatory networks controlling highly conserved wing patterning genes (Wallbank et al. 2016; McMillan et al. 2020). Variation in enhancers can also explain phenotypic differences between individuals of the same species. Pelvic loss in populations of threespine stickleback occurred through modifications to a tissue-specific enhancer of *Pitx1*, and positive selection on this enhancer is associated with the reduction of the pelvis in these populations (Chan et al. 2009). Coat color variation in the oldfield mouse (*Peromyscus polionotus*) is associated with an enhancer region upstream of the *Agouti-signaling protein* coding region (Wooldridge et al. 2022). These examples, among many others, demonstrate that changes in enhancers are often a major source of phenotypic variation, at least for morphological traits (Carroll 2000; Prud’homme et al. 2007).

To explore the contribution of enhancers to social evolution, we identified and quantified the activity of regulatory regions genome-wide in *L. albipes* using the high-throughput enhancer discovery tool, STARR-seq (self-transcribing active regulatory region sequencing, (Arnold et al. 2013)). We compared the activity of enhancers across social and solitary populations of this socially flexible bee to identify enhancer regions associated with social variation. Combining measures of enhancer activity with allele frequency variation within these regions, we identified putative causal variants underlying enhancer activity variation. Finally, we leveraged population genetics data to test whether differences in the divergence rates of genetic variants are associated with enhancer activity. Taken together, this study provides a comprehensive overview of enhancer variation and evolution within a socially flexible bee.

## Results

### Upgraded *L. albipes* genome assembly

We used a combination of PacBio, HiC, and HiFi sequencing technologies to build an upgraded *L. albipes* assembly for use in characterizing enhancer elements. Circular consensus sequencing of a PacBio SMRTBell library resulted in a yield of 22.92 Gb across 1,963,023 HiFi reads with an N50 read length of 11.67 kb. Of those reads, 835 (0.04%) HiFi reads contained artifact adapter sequences and were discarded. The remaining 1,962,188 HiFi reads (99.96% of the total) were used for contig assembly and represented 50x coverage of the genome. Short-read sequencing of the HiC library resulted in 122,133,276 read pairs which exceeds the 100 million read pairs for 1 Gb of genome recommended for HiC data and represents 53x coverage of the genome.

We assembled the genome into a total of 68 contigs, all of which were placed into 25 chromosomes and the mitochondrial genome as one contig with a total size of 344.23 Mb (Table S1, Fig. S1-S2). Estimation of consensus accuracy of the assembly relative to the data used to generate the assembly revealed a raw quality value (QV) of 54.973 and an adjusted quality value of 55.864, which shows that the assembly is an accurate representation of the reads sequenced.

### Characterization of enhancers across 6 populations of *Lasioglossum albipes*

We generated STARR-seq (Arnold et al., 2013) libraries to identify and quantify enhancer activity across six populations of the socially flexible bee, *L. albipes* (Fig. 1a). For each population, we transfected 3 independent flasks of cells which were highly correlated in both genomic coverage of input libraries and STARR-seq plasmid derived mRNA (average correlations of 92.8% and 90.3% within populations, respectively) (Fig. S3). 36,216 regions of the *L. abipes* genome showed significant enhancer activity (Fig. 1b; Table S2). Enhancer regions were variable in size but averaged 1278 nucleotides in length (1277.67+/-691.75 sd; Table S2), which is near the upper end of most estimates of enhancer sizes across insects and vertebrates (Levine and Tjian 2003; Whyte et al. 2013; Panigrahi and O’Malley 2021). Importantly, because our average fragment length of tested regions was 504 bp (+/-96 bp), this creates a lower bound for measuring active regions (i.e., a small enhancer element contained within a 500bp fragment will lead to amplification of the entire fragment). Therefore, our map of ∼36k regions includes both enhancers as well as flanking regions, and the true enhancer length is likely closer to published estimates of anywhere from 10-1000bp. Our identified enhancer regions were located genome-wide (Fig. S4) and were especially prevalent within introns (34%) or in intergenic regions (i.e., more than 10kb from any annotated gene; 38%).

**Figure 1.**
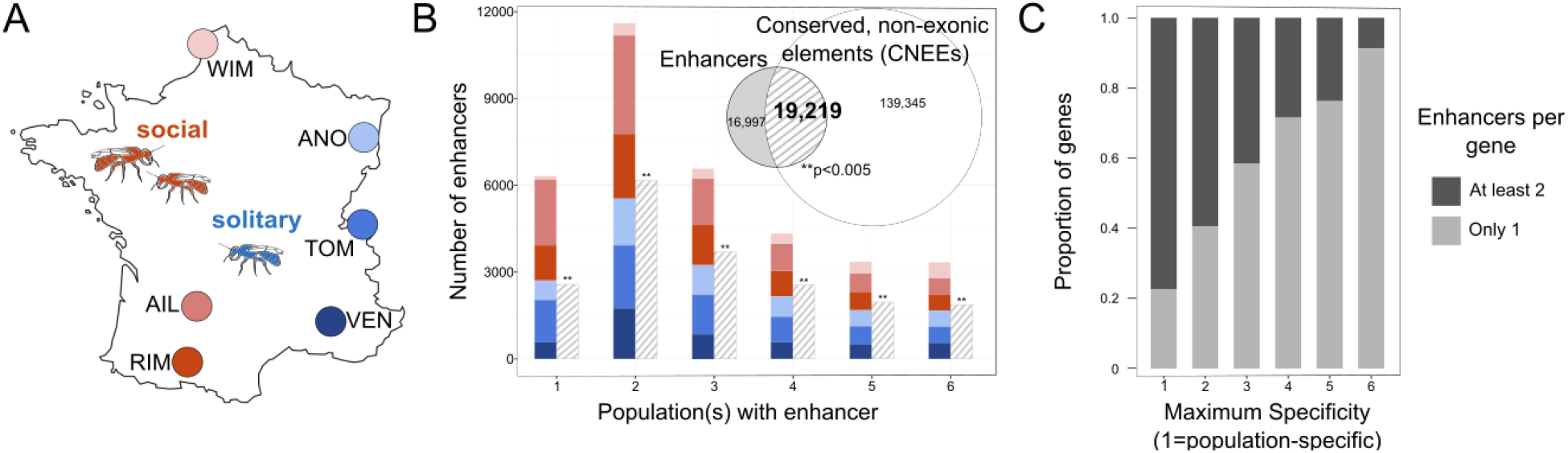
The *Lasioglossum albipes* genome contains both conserved and population-specific enhancers. (A) Location of six populations in France sampled for STARR-seq characterization of genome-wide enhancer activity. Western populations (WIM, AIL, RIM) express social behavior while eastern populations (ANO, TOM, VEN) express solitary behavior through independent, convergent losses of sociality (Kocher et al. 2018). (B) Distribution of enhancers identified in different numbers of populations. Solid bars are colored with the proportion of enhancers within a bar active in each population. Hashed bars show the number of enhancers in each set that directly overlap with bioinformatically-predicted regulatory regions (Conserved, Non-Exonic Elements, CNEEs) in a previous study (Jones et al. 2023). Enhancers overlap CNEEs more often than expected by chance across all categories shown, including the set of all enhancers shown in the Venn diagram inset (**p<0.005, permutation tests using regioneR package of R and randomly shuffled regions). (C) Genes near population-specific enhancers (maximum specificity=1) are more likely to also be proximal to additional enhancers. A maximum specificity of 6 indicates that all enhancers assigned to a given gene are active in all six populations (i.e., conserved).

Nearly 90% of annotated genes (9865/10979) were within 10kb of at least one enhancer. Across enhancers, 4,029 (∼11%) were active in all 6 populations, 18,928 (50.5%) were active in at least 3 populations, and 6,317 were unique to just one population (Fig. 1b). Of genes assigned to these population-specific enhancers, 77% were also predicted targets (based on strict priority assignment, see Methods) of at least one additional enhancer. Genes proximal to enhancers active in greater numbers of populations were less likely to also be assigned to additional enhancers (Fig. 1c). Fully conserved enhancers (i.e., enhancers active in all 6 populations) were proximal to a set of genes enriched for many (87) GO terms, including terms related to chemotaxis, synapse organization, morphogenesis, and regulation of cellular and organismal processes (Table S3). These results suggest that core molecular functions may be maintained through highly conserved enhancers, while other traits are regulated by enhancers that are more evolutionarily labile.

Despite the extensive turnover of enhancers among *L. albipes* populations, we found evidence of possible compensation for enhancer losses. Lineage-specific enhancers were repeatedly proximal to the same genes; 54 of 57 combinations of two or more lists of population-specific enhancer targets overlapped more than expected by chance (Table S4) (Wang et al. 2015). These results are consistent with previous findings across *Drosophila species* (Arnold et al. 2014), where compensatory enhancers evolved in lineages where ancestral enhancers were lost, maintaining gene activity of the target regions.

Previous comparative genomic analysis of bees identified a set of loci predicted to have regulatory function based on sequence conservation across species (Jones et al. 2023). We compared the location of our STARR-seq enhancers with these conserved, non-exonic elements (CNEEs) in *L. albipes* and found that 53% of all enhancers overlapped at least one CNEE, 1.27x more overlap than expected by chance (permutation test, z-score=45.8283, p<0.005). Enhancers significantly overlapped CNEEs regardless of the number of populations in which they were active (Fig. 1b). These results demonstrate the utility of sequence conservation measures in the identification of active regulatory elements and support the use of STARR-seq in identifying regulatory regions of bees. In addition, these results suggest that enhancers can be conserved across divergent lineages despite the elevated evolutionary rates of enhancers compared with other genomic regions (Villar et al. 2015).

### Differential enhancer activity between social and solitary populations

In addition to characterizing enhancers as present or absent within and across populations, we compared the quantitative activity of enhancers between social and solitary populations. Importantly, solitary populations of *L. albipes* represent independent evolutionary losses of social behavior (Plateaux-Quenu 1993; Plateaux-Quénu et al. 2000; Kocher et al. 2018), enabling us to ask questions about convergence of enhancer activity associated with sociality. Overall, 1182 enhancers (∼3.3% of all enhancers) were significantly different in activity (q<0.05) between social and solitary populations (Fig. 2a, Table S5). 547 differentially active enhancers (DAEs) displayed social-biased activity while 635 DAEs had solitary-biased activity, with no apparent regional clustering in the genome (Fig. S5) or differences in feature location relative to all enhancers (Fig. 2b). Genes proximal to DAEs (i.e., within 10kb) were enriched for many GO terms, including *negative regulation of epithelial cell differentiation* (8.5-fold enrichment), *excitatory synapse* (4-fold enrichment, FDR=0.034), *eye morphogenesis* (3.3-fold enrichment, FDR=0.00005), and *sensory organ morphogenesis* (3.05-fold enrichment, FDR=0.002) (Table S6, Fig. S6). Over one quarter of DAEs (311, 26.3%) were among enhancers present in all six populations (i.e., conserved), compared with 11.1% of all enhancers. These conserved DAEs were enriched for 68 GO terms which clustered by semantic similarity into approximately four groups (Fig. S7; Table S6): cellular and organismal regulation, reproductive and anatomical structure development, behavior and multicellular processes, and cell-cell adhesion. The two GO terms with strongest enrichment were *regulation of toll-like receptor signaling pathway*, with conserved DAEs proximal to all three genes in the *L. albipes* genome annotated with that GO term (54-fold enrichment, FDR=0.007), and *imaginal disc-derived female genitalia development* (33-fold enrichment, FDR=0.024).

**Figure 2.**
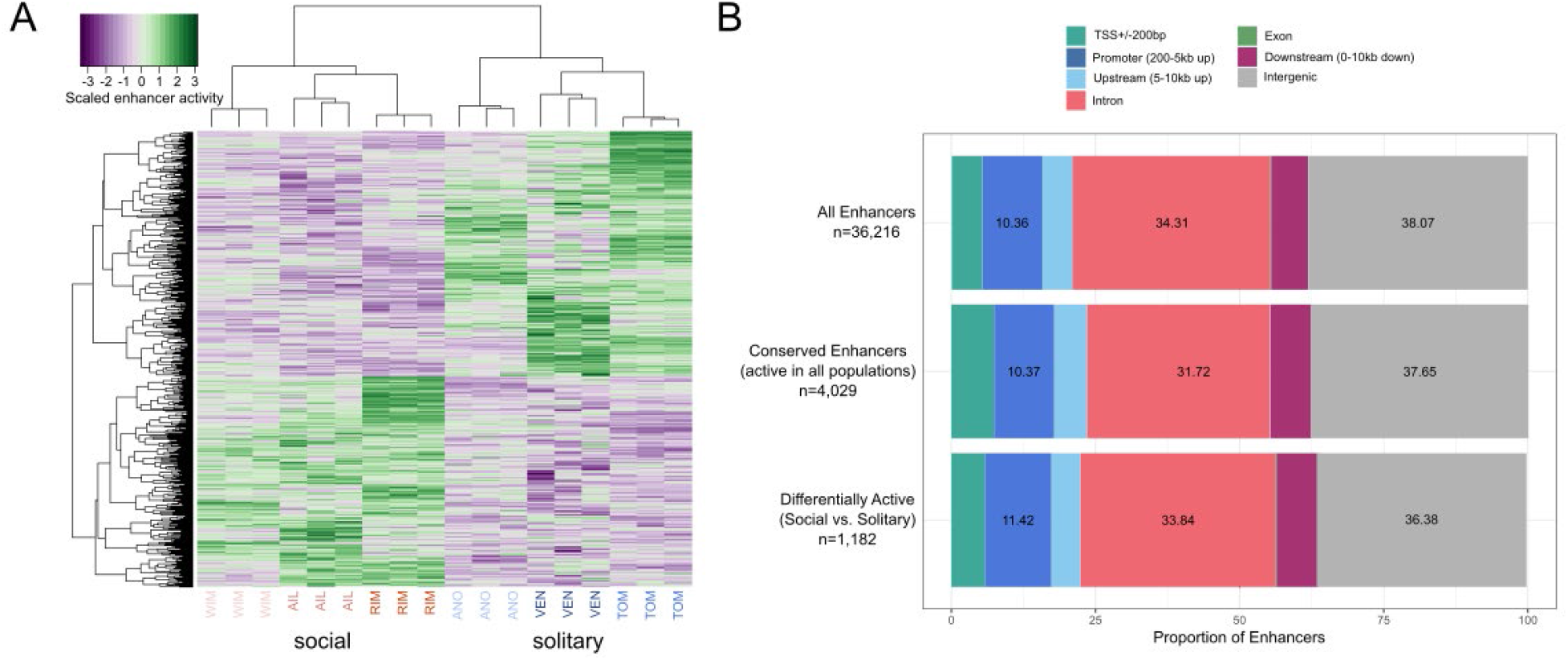
Independent losses of sociality involve convergent changes in enhancer activity. (A) Heatmap of scaled enhancer activity (green=high activity, purple=low activity) for differentially active enhancers (DAEs). Each column is an individual STARR-seq replicate, with three replicates per population, while each row is one of 1182 DAEs. (B) Proportions of enhancers overlapping different gene features, including TSS flanks (TSS+/-200bp), Promoter regions (200bp-5 kb upstream of a gene), Upstream regions (5-10kb upstream of a gene), Introns, Exons, Downstream regions (<=10kb downstream of a gene), and Intergenic regions (no overlap with above gene features).

In addition to frequent conservation of differential enhancers among populations within *L. albipes*, over half of DAEs overlapped conserved regulatory sequences across lineages of halictids. Of the 1182 DAEs, 614 overlapped CNEEs, a proportion similar to the overall rate of CNEE overlap across all enhancers (Fig. 1b; RF=1.23, p<0.005 for overlap between DAEs and CNEEs). These results indicate that, in addition to lineage-specific regulatory changes, social variation in *L. albipes* is also driven by fine-tuning of ancient regulatory elements present across sweat bee species.

Although many enhancers show convergent changes in activity between social and solitary populations (Fig. 2a), population-specific changes associated with the independent losses of sociality may also occur at independent loci regulating similar genes or pathways. In support of this hypothesis, we identified 21 genes targeted by enhancers in all three social, but zero solitary, populations (Table S7). Of those, only eight genes had nearby enhancers with activity in all three social populations, while the remaining 13 genes were proximal to enhancers that were active in only one or two populations each. We additionally identified 52 genes with convergent signatures in solitary populations; 28 of these genes were proximal to a single enhancer that was active in all solitary (but no social) populations. The other 24 genes had at least one proximal enhancer in every solitary population (no single enhancer was active across all solitary populations), and with no assigned enhancers in social populations (Table S7). Notably, only 7 of the 73 genes were significant with the DAE analysis above, highlighting that independent losses of sociality may often involve changes in independent regulatory loci, even if those loci regulate the same genes.

Further evidence of a mix and match between fine-tuning of existing enhancers and emergence of novel regulatory regions comes from a comparison of enhancers to CNEEs faster evolving in either social or solitary lineages across sweat bee species (i.e., reflecting either positive or relaxed selection associated with sociality) (Jones et al. 2023). In general, *L. albipes* enhancers overlap fast-evolving CNEEs more often than expected by chance, (RF=1.36, p<0.005), but DAEs are not more likely to directly overlap fast-evolving CNEEs (p=0.25). At the gene level, however, DAEs and fast-evolving CNEEs are more likely than expected to be proximal to the same genes (341 genes with both a DAE and fast-evolving CNEE; Fig. 3; 1.5-fold enrichment, p=7.49e-20 hypergeometric test of overlap).

**Figure 3.**
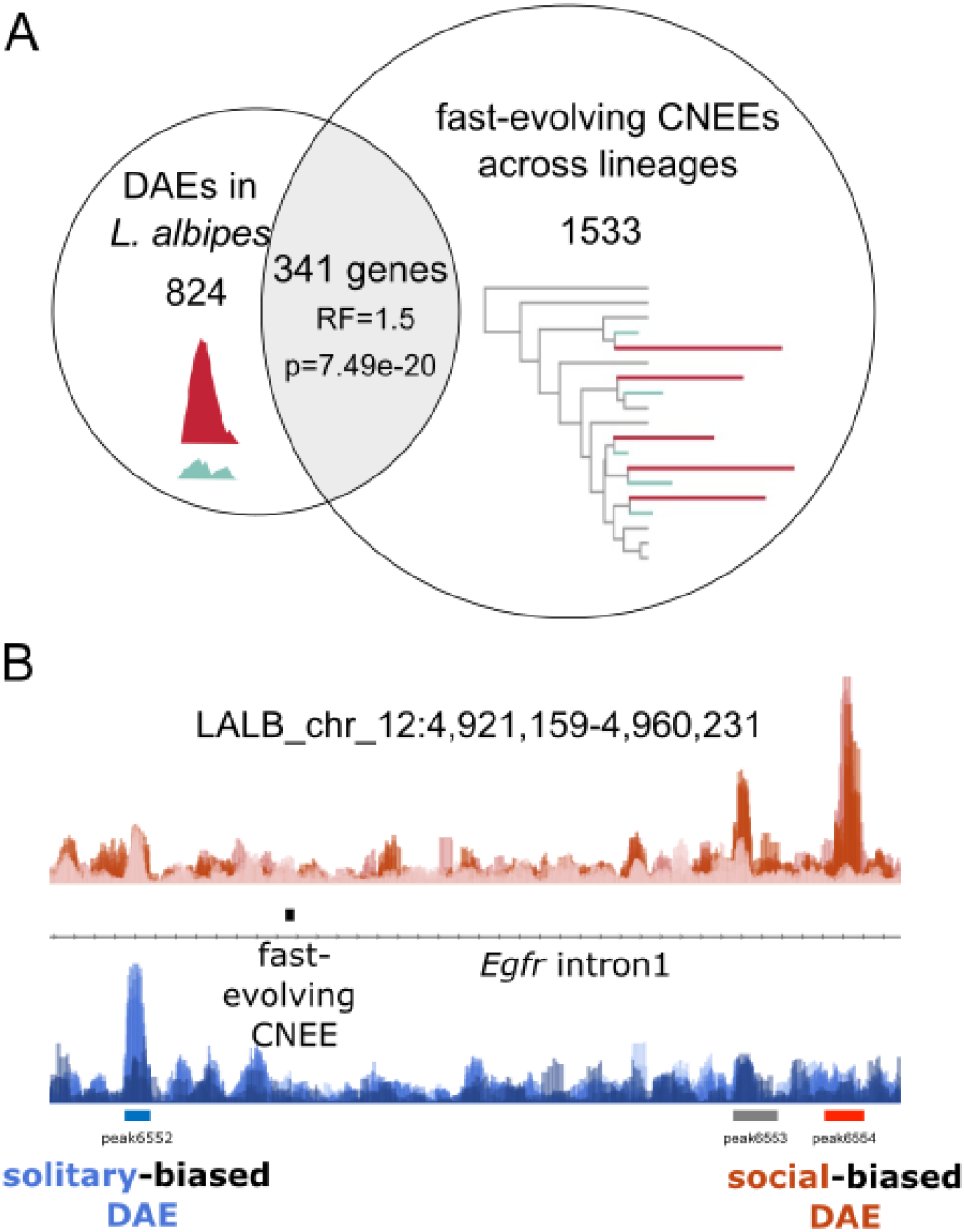
Genes near *L. albipes* enhancers with differential activity between social and solitary populations are also more likely to be near rapidly evolving regulatory regions associated with sociality across species. (A) Overlap of genes proximal to DAEs (left circle, 1165 genes) with genes proximal to CNEEs evolving faster in either social or solitary lineages (1874 genes) (Jones et al. 2023). (B) Example of locus (first intron of *Egfr*) with *L. albipes* enhancers differentially active between social and solitary populations (DAEs) and a fast-evolving CNEE, in this case with longer branch lengths in lineages which have lost social behavior (Jones et al. 2023).

Overlap of DAE-proximal genes and fast-evolving CNEEs was significant for both social-biased DAEs (185 genes, RF=1.8, p=4.11e-17) and solitary-biased DAEs (195 genes, RF=1.5, p=1.42e-10), and overlapping genes were enriched for 170 GO terms (Table S6, Fig. S8). This suggests that both existing and novel regulatory changes can evolve within and among lineages to regulate a shared set of genes associated with social evolution.

To determine whether specific regulatory motifs are repeatedly involved in social evolutionary transitions, we used HOMER (Heinz et al. 2010) and the JASPAR2024 database of insect motifs (Rauluseviciute et al. 2024) to identify enriched TF motifs within DAEs between social and solitary populations. Fourteen motifs were enriched (q<0.05) among social-biased DAEs, while 10 were enriched among solitary-biased DAEs. Six motifs were enriched in both sets (Jra, kay, GATAd, GATAe, srp, and grn), while eight and four were unique to social– or solitary-biased DAEs, respectively (Table S8). Among the social-biased motifs were sequences targeted by proteins that regulate lipogenesis (SREBP), female reproductive gland development and ovulation (Hr39), cell type differentiation of sensory organs (sv), and cholinergic cell fate and T1 neuron morphogenesis in the optic lobe, brain, and CNS (Ets65A) (Thurmond et al. 2019). Solitary-biased unique motifs included Atf3, which is bound by a protein involved in abdominal morphogenesis, the immune system, and metabolic homeostasis (Thurmond et al. 2019), luna, a TF required for proper chromatid separation during cell differentiation, and Atf6, which has numerous functions in regulating gene expression.

### Identification of variants associated with enhancer activity

Because we used genomic DNA fragments isolated from transfected cells as controls, we were able to identify genetic variants within these input libraries and test for correlations between allele frequencies of individual loci and enhancer activity of the genomic fragment containing those loci. Our variant calling pipeline identified 342,829 single nucleotide polymorphisms (SNPs) located within enhancers. Then, using an eQTL statistical framework, we identified significant correlations between the allele frequency of SNPs and enhancer activity for 4071 enhancers, 340 of which are DAEs (Table S9). We assessed the potential impacts of enhancer-correlated SNPs on TF motif binding with the FIMO tool of MEME Suite (Grant et al. 2011). Of enhancers with correlated SNPs, nearly one-third (1311/4071) contained SNPs with significant effects on predicted TF binding (Table S10). The motif with the highest frequency of predicted binding changes was Clamp; allelic variation in 95 enhancer-correlated SNPs was predicted to modify Clamp binding. Notably, this TF was among those identified by HOMER as enriched in social-biased DAEs (Table S8), but only 11 of the 95 SNPs were within DAEs, suggesting variation in Clamp activity may be associated not only with social phenotypes but also with variation in other traits across *L. albipes* populations.

A previous population genomics study of *L. albipes* identified SNPs associated with social behavior across the same populations as in this study (Kocher et al. 2018). We realigned sequencing reads from the previous study to our updated assembly of *L. albipes* and ran a genome-wide, mixed-model association test (GEMMA, (Zhou and Stephens 2012)) to identify variants associated with social behavior following the previously published methods (Kocher et al. 2018). Our reanalysis replicated the results of the original study, with 211 SNPs associated with social behavior, including an intronic SNP of *syx1a* (Table S11; 194 SNPs were identified in the previous study with the same FDR-corrected significance threshold) (Kocher et al. 2018). Our sampling of populations for STARR-seq was independent and therefore included only a subset of the same variants tested in the previous study. Comparing all non-pruned variants in the published data with those sampled in this study, we identified 230,328 SNPs located within enhancers and common to both studies (by chance, the intronic *syx1a* SNP was not sampled within the STARR-seq dataset). Of these common variants, 5104 had significant allele frequency correlations with enhancer activity and 34 were significant with our reanalyzed GEMMA analysis (adjusted Wald p-value<5e-5, as in (Kocher et al. 2018)), though no SNPs were significant in both analyses. Using a less conservative threshold for GEMMA (adjusted Wald p<0.05), eight SNPs were both associated with sociality in the population genomic dataset and had allele frequency correlations with enhancer activity (Table S11).

### Enhancer evolution is associated with signatures of selection

Enhancers which are functionally relevant in mediating social traits are expected to be adaptive and therefore show signatures of evolutionary divergence between social and solitary populations. Using previously published population genetic data, we identified variants and calculated the Population Branch Statistic (PBS, (Yi et al. 2010)) for social and solitary populations at each site for 448,477 SNPs within enhancers, using the closely related sister species, *L. calceatum* (also a socially flexible species), as the outgroup. We additionally used *L. calceatum* to infer the ancestral allele for each biallelic SNP. We combined this information with the results of our analysis of enhancer-correlated SNPs to determine whether derived alleles are associated with an increase or decrease in enhancer activity. Consistent with a pattern of positive selection on functional sites within enhancer sequences, derived alleles associated with an increase in enhancer activity in both social– and solitary-biased DAEs exhibited longer PBS branch lengths (Fig. 4a-b). In contrast, divergent sites within non-DAEs were more likely to lead to decreases in enhancer activity in social populations (Fig. 4a).

**Figure 4.**
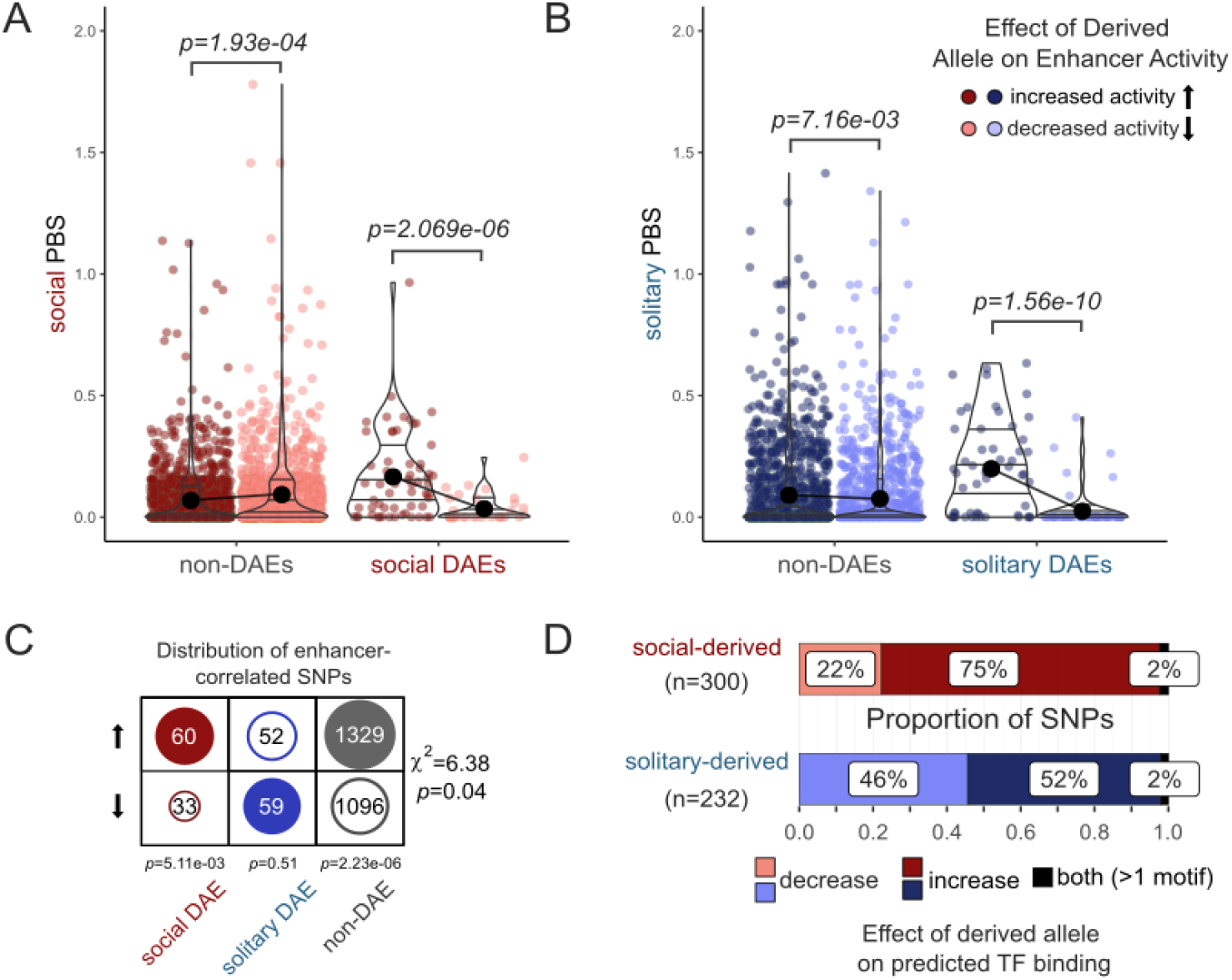
New mutations that influence enhancer activity show signatures of selection in both social and solitary populations. **(A)** Population branch statistic (PBS) values for social populations are elevated for sites where the derived allele leads to increased activity (dark red) of enhancers in social-biased DAEs, while the opposite pattern is observed in non-DAEs (i.e., sites within non-DAEs tend to have longer PBS values if the derived allele leads to a decrease in enhancer activity). p-values from Kruskal-Wallis Rank Sum Test. **(B)** PBS values for solitary populations are elevated for both non-DAEs and solitary-biased DAEs for sites where the derived allele leads to an increase (dark blue) in enhancer activity compared with sites where the derived allele leads to a decrease (light blue) in enhancer activity. p-values from Kruskal-Wallis Rank Sum Test. **(C)** Social-biased DAEs are more likely to contain SNPs where the derived allele leads to an increase in enhancer activity compared with solitary-biased DAEs. Non-DAE sequences are similarly more likely to contain sites where derived alleles lead to increases in enhancer activity, while SNPs within solitary-biased DAEs were equally likely to contain new mutations that lead to increases or decreases in enhancer activity. Pearson’s Chi-squared=6.38, p=0.04, *p-*values below each column are from one-sample binomial tests. **(D)** Social-derived alleles are more likely to lead to increases in predicted TF binding (p=2.40e-56, proportion test). Direction of effect is based on comparison of FIMO-predicted binding scores for alleles of SNPs correlated with enhancer activity across populations. All significant binding effects are provided in Table S12.

Among social-biased DAEs and non-DAEs, we also found that derived alleles are more likely to lead to increased enhancer activity than expected by chance (Fig. 4c). Variants in solitary-biased DAEs did not show this same pattern; these enhancers contained similar proportions of derived alleles which increased or decreased enhancer activity. In addition, 75% of social-derived alleles with predicted effects on TF binding led to increases in binding scores, while solitary-derived alleles affected TF binding in both directions (Fig. 4d). Together, these results are consistent with selection on causal variants within enhancer regions of *L. albipes*, and specifically suggest that alleles leading to increased regulatory activity in social populations experience directional selection. In populations which have secondarily reverted to solitary life history strategies, on the other hand, relaxed selection on enhancers ancestrally involved in social behavior may lead to the accumulation of alleles that reduce activity of these regions. This pattern is consistent with previous reports of relaxed selection on both coding and non-coding regions among lineages that have lost social behavior (Jones et al. 2023). Our results reinforce these findings and suggest that turnover of regulatory regions may be playing an underappreciated role in shaping behavioral traits even among closely related populations.

## Discussion

In this study, we took advantage of the convergent losses of social behavior in a socially flexible bee, *Lasioglossum albipes*, to ask whether and how changes in enhancer function may be linked to social evolution. We adapted a high-throughput enhancer reporter assay to measure the activity of gene regulatory regions in six populations of *L. albipes*, representing three independent losses of social behavior. We found active enhancers genome-wide by testing fragments of *L. albipes* genomic DNA in a reporter construct transfected into *Drosophila* S2 cells. Our study demonstrates the ability to use the cellular machinery of a model species (*Drosophila*) to measure the activity of gene regulatory elements of a different organism. We were specifically interested in differences in enhancer activity related to social phenotypes, but further research could examine the role of enhancer activity in other traits in these bees using our genome-wide enhancer maps as a starting point.

Our identified enhancers overlapped significantly with bioinformatically predicted regulatory regions of bees (Jones et al. 2023), suggesting *Drosophila* cell lines and their associated proteins can indeed activate bee enhancer elements. This is not surprising given the high conservation of TFs and their binding specificities. TFs and the DNA motifs they bind to are almost unchanged across Bilateria, with flies and humans sharing nearly the same gene regulatory code despite over 600 million years of evolution (Nitta et al. 2015). A subset of the enhancers we characterized were active in all six of the populations we examined. These enhancers were proximal to genes enriched for GO terms related to chemotaxis, synapse organization, and morphogenesis, suggesting that such highly conserved enhancers may help to maintain core molecular processes.

While TFs and their binding motifs are highly conserved, enhancer elements themselves can display rapid evolutionary turnover. For example, the split between *D. melanogaster* and *D. yakuba* involved the gain of hundreds of enhancer elements, the majority of which arose *de novo* from non-functional sequences (Arnold et al. 2014). We discovered that even among closely related populations, enhancers are gained and lost at an impressive rate. Over 6,000 enhancers were active in only one population, pointing to substantial intraspecific genetic variation in *L. albipes* affecting gene regulatory activity. Similar turnover has also been observed among populations of *Heliconius* butterflies (Lewis and Reed 2019). Interestingly, the *L. albipes* population-specific enhancers tended to be proximal to an overlapping set of genes, suggesting that compensatory enhancers may arise when ancestral enhancers are lost.

We identified a subset of enhancers that had differential activity between social and solitary *L. albipes* populations (DAEs). These enhancers were near genes with functions related to neuronal development, including multiple enriched GO terms related to axon recognition and neuron projection, as well as photoreceptor development. While these processes play generalized and fundamental roles in neurodevelopment, it is intriguing to note that previous work in *L. albipes* identified differential investment in olfactory sensilla between social and solitary populations (Wittwer et al. 2017). Axon targeting is especially important during the development of the olfactory system in insects (Komiyama and Luo 2006), including in the projection of olfactory receptor neurons into antennal glomeruli and the neurons that integrate olfactory information in the mushroom bodies (Opachaloemphan et al. 2018). Moreover, the TF binding motif for shaven (sv), was enriched in social-biased DAEs. Shaven is a TF associated with sensory organ morphogenesis and antennal development (Fu et al. 1998; Scalzotto et al. 2022). Together, these results suggest that differences in sensory perception between social and solitary populations of *L. albipes* may be mediated in part through differences in the activity of enhancers regulating genes involved in central nervous system and sensory organ development.

DAEs associated with behavior were more than twice as likely to be active in all six populations compared to all characterized enhancers, and half of these DAEs also overlapped evolutionarily conserved CNEEs identified across different sweat bee species (Jones *et al* 2023). In contrast, we also identified population-specific changes associated with the independent losses of eusociality in *L. albipes*. We identified a subset of genes with proximal enhancers in only social (n=21) or only solitary (n=52) populations, suggesting that different enhancers may be independently gained or lost across lineages in a compensatory manner to regulate a shared set of genes relevant to social behavior. Furthermore, DAEs are not more likely than chance to overlap CNEEs that are fast-evolving across species, but they are more likely to be proximal to the same genes. Together, these results imply that variation in social behavior is most likely mediated by a combination of the fine-tuning of ancient regulatory regions as well as the emergence of novel, lineage-specific regulatory elements.

Similar to previous studies of social evolution across distantly-related bee lineages (Kapheim et al. 2015; Rubin et al. 2019; Jones et al. 2023), we find evidence that selection on gene regulatory elements plays a role in social transitions within *L. albipes*. Enhancers with differential activity between social and solitary populations contained variants with longer branch lengths compared with those in enhancers with conserved activity, hinting at either positive directional or relaxed purifying selection at these loci.

Relaxed selection on protein coding genes has been identified as a possible driver of social insect caste evolution (Hunt et al. 2011) and in the evolution of immunity genes in the honey bee, *Apis mellifera* (Harpur and Zayed 2013). Relaxed selection is also prevalent among protein coding genes in lineages which have reverted from social to solitary life history strategies in bees (Jones et al. 2023) and in social spiders (Tong et al. 2022). We expected to see a similar pattern of relaxed selection in non-coding regions of the genome, and while we did identify SNPs within enhancers that had long branch lengths in solitary populations, many derived variants in these regions led to an increase, rather than decrease, in overall enhancer activity and predicted motif binding. These results are consistent with positive selection on loci that increase regulatory function of these regions, a pattern which we identified in both social– and solitary-biased enhancers. Positive selection on enhancer regions has been reported for certain classes of enhancers, including those regulating immune-related functions in humans (Moon et al. 2019) and a rapidly evolving enhancer conferring gain-of-function for human-specific limb expression (Prabhakar et al. 2008). Interestingly, we also identified many enhancers containing variants diverging more rapidly in social populations that are associated with reductions in enhancer activity. These loci may reflect the fine-tuning of gene expression patterns required to produce both queen and worker phenotypes from a shared genome, or they may reflect modifications to enhancers that are increasingly important in solitary populations of *L. albipes* but experiencing drift in social populations.

Solitary populations not only retain much of the enhancer repertoire of social ancestors but also gain activity in many regulatory regions, which is somewhat unexpected. Multiple studies of bee social evolution have identified an expansion of TF motifs in the genomes of social bees (Kapheim et al. 2015; Jones et al. 2023), with a secondary reduction in motif presence when sociality is lost (Jones et al. 2023). However, these studies rely upon bioinformatic predictions of regulatory regions and are limited to studying proximal regulatory regions of highly conserved and non-duplicated genes to enable cross-species comparisons. Indeed, 38% of enhancers we identified in *L. albipes* were more than 10kb from any annotated genes, including many enhancers with differential activity between social and solitary populations. Future studies integrating these data with functional examinations of chromatin accessibility and gene expression across different developmental timepoints and tissues are needed to further resolve these observations.

## Conclusions

Our findings suggest that gene regulatory variation is indeed a contributor to social variation in *L. albipes*, and that selection on enhancer elements may facilitate transitions in eusociality. We found evidence that conservation of enhancers may be associated with core molecular processes, and that compensatory enhancers can evolve among independent lineages, potentially to help maintain regulatory activity of target genes in the face of high levels of enhancer turnover. A subset of enhancers showed behavior-specific activity patterns across social forms. Our results reveal that both existing and novel regulatory changes can evolve to regulate a shared set of genes associated with social evolution. Taken together, our results indicate that variation in social behavior is driven both by fine-tuning of ancient regulatory elements as well as by lineage-specific regulatory changes.

## Methods

### Upgraded *L. albipes* genome assembly

From two male *L. albipes* bees, we prepared a PacBio SMRTbell library and a short-read HiC library which were sequenced on a PacBio Sequel IIe and an Aviti Element 2×150 flow cell, respectively. After sequencing and circular consensus analysis using SMRTLink v11.0.0.144466, we identified adapter-contaminated HiFi reads and omitted them from the HiFi read pool using HiFiAdapterFilt v2.0 (Sim et al. 2022) and used the remaining HiFi reads to assemble the *L. albipes* genome into contigs using HiFiASM v0.15.1-r328 (Cheng et al. 2021). After contig assembly, we used the YAHS v1.1 (Zhou et al. 2023) pipeline to create a HiC contact map (Fig. S1) using the 2×150 paired HiC reads and visualized and manually edited the HiC contact map using Juicebox v1.11 (Durand et al. 2016). After HiC scaffolding, non-arthropod contigs were omitted from the final assembly after taxonomic identification using BLAST+ v2.13.0+ (Camacho et al. 2009) to the NCBI nucleotide database (accessed: 2022-02-14) and using DIAMOND v2.0.9 (Buchfink et al. 2021) to the UniProt database (accessed: 2022-06-01), the results of which were summarized using blobtools2 v.4.1.5 (Challis et al. 2020; Sim, 2022: https://github.com/sheinasim/blobblurb). The final assembly was estimated for base accuracy relative to the HiFi reads using the program YAK which is a part of the HiFiASM pipeline (Cheng et al. 2021). Raw data and assembly are deposited on NCBI’s SRA with accession IDs SUB14643391 and SUB14646709, respectively. Note that the NCBI version of the assembly includes all contigs, so we have also uploaded the filtered assembly (removing all non-arthropod and unplaceable no-hits) used in our analyses on our github project page: https://github.com/kocherlab/Lalbipes_STARRseq/.

### Genome annotation

We created a snakemake pipeline to annotate the PacBio assembly using BRAKER3 (Gabriel et al. 2024). BRAKER3 requires a soft-masked assembly, a BAM of merged RNAseq samples, and a database of protein sequences to provide homology information. We masked the assembly using a combination of RepeatModeler (v2.0.4) (Flynn et al. 2020) and RepeatMasker (v4.1.4) (Smit et al., 2008-2015), then created a soft-masked assembly using the python script softmask.py. We aligned 40 paired-end *Lasioglossum albipes* RNAseq samples (NCBI SRA accession PRJNA1142947; distinct samples from the STARR-seq libraries above) to the assembly using HISAT2 (v 2.2.1) (Kim et al. 2019), then sorted and merged them using SAMtools (v 1.16.1) (Danecek et al. 2021). We created a protein sequence database using *Apis mellifera* (AMEL_HAv3.1), *Bombus impatiens* (BIMP_2.2), ortholog groups with an arthropod ortholog from OMA (Altenhoff et al. 2024), and the arthropods database from OrthoDB (odb11; (Zdobnov et al. 2021)). All software was run using default parameters. The annotations resulted in 10,979 genes with a BUSCO score (Simão et al. 2015) of 97.7% using the Hymenoptera odb10 database. GFF3 file for annotation is available on the project github: https://github.com/kocherlab/Lalbipes_STARRseq/.

### Generation of STARR-seq plasmid libraries

Genomic DNA was isolated from individuals collected from each of the six *L. albipes* populations using the *Quick-*DNA Microprep Plus Kit (Zymo Research cat. no. D4074). Details on individuals and tissues sampled for each population are in Table S12. A pool of 5 μg from each population was sonicated in a microTUBE with AFA fiber (cat. no. 520045) using a Covaris LE220 and the following parameters: 45 sec sonication time, 450W peak incident power, 15% duty factor, 200 cycles per burst. Sheared DNA was run on a 1% agarose gel and 450-750bp fragments were size-selected via excision under blue light. Size-selected DNA was purified using a Gel DNA Recovery Kit (Zymo Research cat. no. D4008), followed by an additional purification with the QIAquick PCR Purification Kit (Qiagen cat. no. 28104). Size-selected DNA was quantified using a Qubit dsDNA HS Assay Kit (Invitrogen cat. no. Q32854) on a Qubit4 Fluorometer.

Illumina-compatible adapters were ligated to size-selected genomic DNA using the NEBNext Ultra II End Repair Module (NEB E7546L) with 1 μg fragmented DNA per population. Adapter-ligated DNA libraries were then cleaned with a 1.8x volume ratio of AMPure XP Reagent beads (Beckman Coulter cat. no. A63881) to sample following manufacturer protocols for PCR Purification, then cleaned a second time with 0.8x bead to sample volume ratio. Adapter-ligated DNA libraries were then amplified in 5 separate reactions for 5 cycles (PCR conditions: initial denaturation of 98C for 45 sec, then 5 cycles of 1) denaturation: 98C for 15 sec, 2) annealing: 65C for 30 sec, 3) elongation: 72C for 45 sec, and a final elongation at 72C for 60 sec) each using in_fusion_F (TAGAGCATGCACCGGACACTCTTTCCCTACACGACGCTCTTCCGATCT) and in_fusion_R (GGCCGAATTCGTCGAGTGACTGGAGTTCAGACGTGTGCTCTTCCGATCT) primers at 10 uM and 2x KAPA HiFi HotStart Ready Mix (Roche cat. no. KK2601). PCR reactions were pooled for each population and cleaned with 0.8x bead to sample volume ratio with AMPure XP Reagent beads (Beckman Coulter cat. no. A63881) followed by an additional purification using the QIAquick PCR Purification Kit (Qiagen cat. no. 28104). Libraries were quantified using a Qubit dsDNA HS Assay Kit (Invitrogen cat. no. Q32854) on a Qubit4 Fluorometer and average sizes of each adapter-ligated, amplified library was determined with Agilent High Sensitivity DNA reagents on an Agilent 4200 TapeStation (Agilent cat. Nos. 5067-5592, 5067-5593, 5067-5594).

pSTARR-seq_fly was a gift from Alexander Stark (Addgene plasmid #71499; http://n2t.net/addgene:71499; RRID:Addgene_71499) (Arnold et al., 2013). The pSTARR-seq_fly reporter vector was digested with AgeI-HF (NEB cat. no. R3552S) and SalI-HF (NEB cat. no. R3138S) restriction enzymes with 250 units of each and 25 μg vector per reaction, incubated at 37C for 2h followed by a 20 min heat inactivation at 65C. Digested products were run on a 1% agarose gel and linearized vector was selected via excision under blue light and purified using a Gel DNA Recovery Kit (Zymo Research cat. no. D4008). Eluates from gel extraction were purified using a 1.8x bead cleanup with AMPure XP beads (Beckman Coulter cat. no. A63881). Adapter-ligated DNA libraries were cloned into purified pSTARR-seq_fly using a 2:1 molar ratio of insert (size determined via TapeStation) to plasmid (∼4125bp). Two cloning reactions were conducted for each population using 1 μg digested plasmid, the appropriate amount of PCR amplified, adapter-ligated DNA library to have a 2x molar excess insert to plasmid, and 10 μl 5x In-Fusion HD Enzyme Premix (Takara cat. no. 638910) in a total volume of 50 μl. Reactions were incubated for 15 min at 50C, then 200 μl EB was added. Next, 25 ul 3M NaAc (pH 5.2) and 2 μl Pellet Paint Co-precipitant (Millipore cat. no. 69049), were added to each reaction, vortexing between each addition. Finally, 750 μl ice-cold 100% EtOH was added, samples were vortexed, then incubated at –20C for 16h. Samples were centrifuged for 15 min at full speed and 4C, vortexed, centrifuged again for 15 min, then supernatant was carefully aspirated. Cloned DNA pellets were washed 3 times with 750 μl ice-cold 70% EtOH, mixing each time by inversion. Cloned DNA pellets were again centrifuged for 15 min at full speed and 4C, supernatant aspirated, then pellets dried for 30 sec at 37C then further at room temperature until dry. Each pellet was resuspended in 12.5 μl EB and incubated for 3h at –80C prior to transfer to –20C for storage until transformation.

Cloned DNA reactions were transformed into electrocompetent MegaX DH10B cells (ThermoFisher cat. no. C640003) using 150 μl cells for each clone (two clones per population) split across two Gene Pulser Electroporation Cuvettes (0.1 cm gap, Bio-Rad cat. no. 1652089) and the Bio-Rad Gene Pulser Xcell system with the following electroporation conditions: 2 kV, 25 μF, 200 ohm. Immediately after electroporation, 500 μl of pre-warmed recovery medium was added and cells were transferred to round bottom tubes with an additional 4 ml warm recovery media. Transformed bacteria were incubated for 1h at 37C and 225 rpm, then each transformation reaction was added to 300 ml warm LB+ampicillin (100 μg/ml) in 2L flasks and incubated for 12-13h while shaking at 200 rpm at 37C. Cells were harvested via centrifugation and plasmids were purified using the ZymoPure II Plasmid Maxiprep Kit (Zymo Research cat. no. D4203) with a maximum of 75 ml culture per column and eluted in water heated to 50C prior to elution. Plasmids were pooled within each clone and then across clones from the same population, resulting in one clone library per population.

### Transfection of *Drosophila* cells with STARR-seq plasmid libraries

Three replicate flasks were seeded and transfected for each *L. albipes* population. *Drosophila* S2 cells (S2-DRSC; DGRC Stock 181; RRID:CVCL_Z992, (Schneider, 1972)) were cultured in M3 media (Sigma-Aldrich cat. no. S8398) supplemented with BactoPeptone (BD Biosciences #211677), yeast extract (Sigma-Aldrich Y1000), 10% heat-inactivated fetal bovine serum (Gibco cat. no. 10437-010) and 1% Penicillin/Streptomycin solution (ThermoFisher cat. no. 15140122) at 25C using standard cell culturing protocols. Twenty four hours prior to transfection, cells were split, washed, counted, and seeded at a density of 27 million cells in 15 ml media per T75 flask, with 3 flasks seeded per population. Effectene Transfection Reagent (Qiagen cat. no. 301427) was used to transfect 3 replicate flasks of cells per population. For each replicate, 12 μg plasmid clone library was diluted with Buffer EC to a total volume of 450 μl per flask, 96 μl enhancer was added and the reaction was vortexed briefly then incubated for 5 minutes. 150 μl effectene was added, the solution was mixed by pipetting up and down 5 times, then incubated for 10 minutes. One ml media was added to the complex, mixed by pipetting up and down twice, then added drop-wise onto the flask of cells. Flasks were swirled gently to ensure uniform distribution then returned to 25C incubator for 48 hours until harvest.

For harvesting, cells were gently pipetted to bring into suspension then centrifuged for 5 min at 350g. Cells were washed once with 10 ml 1X PBS. then incubated at 37C for 5 min in 2 ml M3+BPYE media containing 1 ml Turbo DNase (2 U/μL) (ThermoFisher cat. no. AM2239) per 36 ml. Cells were again pelleted via centrifugation, supernatant removed, then resuspended 10 ml 1X PBS. An aliquot of 10% unlysed cells per flask were set aside for later plasmid extraction (pelleted and stored at –20C) and the remaining cells were pelleted and lysed in 2 ml RLT (Qiagen RNeasy Midi Kit cat. no. 75144) plus 20 μl 2-mercaptoethanol (Sigma cat. no. 60-24-2) then frozen at – 80C.

### Generation of input and STARR libraries

For each of the eighteen transfected flasks, both an input library (derived from fragment inserts of plasmid DNA purified from transfected cells) and a STARR library (derived from plasmid-derived mRNA) were generated. Plasmid DNA was purified from 10% of harvested cells per flask using a QIAprep Spin Miniprep Kit (Qiagen cat. no. 27104). Total RNA was extracted from 90% of harvested cells per flask using a Qiagen RNeasy Midi Kit (Qiagen cat. no. 75144). mRNA was isolated from 75 μg total RNA using Dynabeads Oligo(dT)_25_ (Invitrogen cat. no. 61005), followed by a DNase digestion with Turbo DnaseI (ThermoFisher cat. no. AM2239). RNA was cleaned with RNAClean XP beads (Beckman Coulter cat. no. A63987) using a 1.8x bead to sample volume ratio and reverse transcribed using SuperScriptIII (Invitrogen cat. no. 18080093) using a gene-specific primer (CTCATCAATGTATCTTATCATGTCTG). cDNA was purified using a 1.8x bead to sample volume of AMPure XP beads (Beckman Coulter cat. no. A63882) and used in a junction PCR to amplify only plasmid-derived mRNA with primers that span a synthetic intron of the pSTARR-seq_fly reporter vector. cDNA was used in this junction PCR with 15 cycles (PCR conditions: initial denaturation of 98C for 45 sec, then 15 cycles of 1) denaturation: 98C for 15 sec, 2) annealing: 65C for 30 sec, 3) elongation: 72C for 70 sec, and a final elongation at 72C for 60 sec) each using junction_F (TCGTGAGGCACTGGGCAG*G*T*G*T*C) and junction_R (CTTATCATGTCTGCTCGA*A*G*C) primers and 2x KAPA HiFi HotStart Ready Mix (Roche cat. no. KK2601). Junction PCR products were purified with a 0.8x bead to sample volume ratio with AMPure XP Reagent beads (Beckman Coulter cat. no. A63881). Either 10 ng plasmid DNA (input) or entire cleaned junction PCR products (STARR libraries) were used as input for a PCR to add indices for sequencing, and final libraries were quantified using the dsDNA HS Assay Kit (Invitrogen cat. no. Q32854) on a Qubit4 Fluorometer and average sizes of each library was determined with Agilent High Sensitivity DNA reagents on an Agilent 4200 TapeStation (Agilent cat. Nos. 5067-5592, 5067-5593, 5067-5594). Libraries were sequenced on 3 flowcells (i.e., 6 lanes) of a NovaSeq SP with 2×50nt paired-end reads at the Genomics Core Facility at Princeton University. Average genome coverage across samples was 104x (range: 78x-140x), with 97.5% and 93.3% of bases covered by ≥10 or ≥20 reads in input samples, respectively (Fig. S9). Additional information on sequencing coverage is in Table S13.

### STARR-seq data processing

Raw FASTQ files from STARR-seq input and plasmid-derived mRNA were processed to remove low quality bases and adapter contamination using fastp (Chen et al. 2018) with default parameters. Processed FASTQ files were then aligned to the *Lasioglossum albipes* genome with bwa mem (Li 2013) and sorted with samtools (Danecek et al. 2021). Information on fragment sizes and mapping rates are in Table S13. Enhancer peaks were called with MACS2 (Zhang et al. 2008) on each replicate flask, with input libraries as controls and the following parameters: –f BAMPE, –g 3.44e8 –keep-dup all –q 0.05. Sequencing reads are available in NCBI’s Sequence Read Archive under BioProject PRJNA980186.

### Identification of enhancers and differential enhancers

Peaks called on each flask (3 per population, 18 total) were concatenated, sorted and merged with BEDtools (Quinlan and Hall 2010). Merged peaks called with MACS2 (Zhang et al. 2008) were filtered to include those detected in a minimum of 2 flasks with a maximum length of 10kb, resulting in a consensus peak set of 36,216 enhancers. This consensus set was used with featureCounts from the Subread package (Liao et al. 2014) to count reads mapping to each peak region for all input and STARR libraries. Assessment of enhancer presence or absence in each population was determined using BEDtools intersect between the consensus set and each MACS2 peak file, and enhancer strength was quantified as the log2 fold-change of normalized counts from mRNA (STARR libraries) relative to DNA input (input libraries) for each replicate.

Differential enhancers between social and solitary populations were identified using the R packages limma and variancePartition and implementation of Dream (Differential expression for REpeAted Measured; (Hoffman and Roussos 2021)). Enhancer regions were tested for significant interactions between social type (social or solitary) and library type (RNA or DNA), with population and sequencing batch as random effects (∼sociality+library+sociality*library+(1|population)+(1|batch)).

### Annotation of proximal genes and motif enrichment analysis

Enhancers were annotated based on proximity to nearest gene models in our updated annotation (see above). When assigning enhancers to features, the following priority was used: tss_flanks (within 50bp of the TSS), tss_upstream (within 200bp upstream of the TSS), promoter (within 5kb upstream of a gene), first intron, intron, 5’ UTR, 3’ UTR, exon, upstream (within 10kb upstream of a gene), downstream (within 10kb downstream of a gene), and intergenic. In the case of a tie (e.g., the enhancer is located within the intron of two different genes), the enhancer was assigned to all genes with the highest priority feature. For “strict” annotation, only the highest priority gene was used, whereas “lenient” annotation includes all genes within 10kb of either side of the enhancer. We used the findMotifsGenome.pl script from HOMER 4.10 (Heinz et al. 2010) to assess the enrichment of insect motifs within enhancer regions using default parameters and the JASPAR2024 database of non-redundant insect motifs (Rauluseviciute et al. 2024). Trinotate v4.0.2 (Bryant et al. 2017) was used to predict gene ontology (GO) terms associated with each gene, and GO enrichment was performed with GOATOOLS (Klopfenstein et al. 2018).

### Overlap with Conserved Non-Exonic Elements (CNEEs)

Overlap of enhancers and Conserved Non-Exonic Elements (CNEEs, (Jones et al., 2023)) was assessed using the RegioneR package in R. Significance of overlap was assessed using the overlapPermTest. Enrichment was calculated as the ratio of observed overlap to the mean of the permuted overlap from overlapPermTest.

### Variant calling on input STARR-seq DNA

We used GATK (v4.3.0.0) (McKenna et al. 2010) to joint-genotype 18 pooled STARR-seq DNA samples from *Lasioglossum albipes*. To prepare the samples for the GATK we trimmed the paired-end FASTQs using fastp (v0.23.2) (Chen et al. 2018) to require reads of at least 30bp after removing adapter content (--detect_adapter_for_pe) and trimming the front and tail. The trimmed FASTQs were then aligned to the assembly using bwa mem (v0.7.17-r1188) (Li 2013) and duplicates were marked using sambamba (v1.0.0) (Tarasov et al. 2015). We used GATK’s HaplotypeCaller to produce GVCFs with the appropriate ploidy for each pooled sample, then generated a VCF from the GVCFs using the GATK’s GenomicsDBImport and GenotypeGVCFs. The VCF was then filtered using the GATK’s VariantFiltration to match the setting used by snpArcher (Mirchandani et al. 2024). Lastly, we filtered the VCF to only include biallelic SNPs that passed a MAF >= 0.05, missing data <= 25%, and QUAL >= 30.

### Identification of alleles correlated with enhancer activity

Allele frequencies were obtained from the input STARR-seq DNA samples per population replicate as above. To test the association between enhancer activity and allele frequencies, the R package MatrixEQTL was used with enhancer activity as the input “expression” matrix. SNPs with no variation were removed prior to running MatrixEQT and p-values were universally corrected for multiple testing with the p.adjust function and method “fdr”. SNPs with significant (FDR<0.05) associations with enhancer activity were then tested for effects on TF motif binding with FIMO (Grant et al. 2011). For each SNP, two 101bp regions (one for each allele) centered on the SNP of interest were scanned with FIMO using the JASPAR2024 database of non-redundant insect motifs (Rauluseviciute et al. 2024). A SNP was considered to have an effect on predicted binding for a given motif if only one allele had a significant (p<0.0001) match for that motif and/or if the ratio of FIMO scores between the two alleles was greater than 1.5, as in (Kapheim et al. 2020).

### Re-analysis of existing population genetic data

We used snpArcher (Mirchandani et al. 2024) to joint-genotype 160 *Lasioglossum albipes* sampled from three social populations (WIM (also referred to as AUD in other publications), AIL (also referred to as DOR), RIM) and three solitary populations (ANO (also referred to as VOS), TOM (also referred to as BRS), VEN) (Kocher et al. 2018). We ran snpArcher using the PacBio assembly with the standard parameters provided in the snpArcher configuration file except for disabling the missingness filter. This was done to allow for greater flexibility when considering missingness in subsequent analyses. We filtered the VCF to only include 139 samples to match those removed in the original analysis in addition to samples with ambiguous records. The 139 samples were then filtered to only include biallelic SNPs that passed a MAF >= 0.05, samples with missing data <= 15, and QUAL >= 30.

### Population Branch Statistic

We computed the population branch statistic (PBS, (Yi et al. 2010)) from Hudson Fst values calculated between *Lasioglossum calceatum*, solitary populations of *L. albipes*, and social populations of *L. albipes* using PLINK (v2.00a3.7LM) (Purcell et al. 2007). The PBS computations for each variant site were performed using calc_pbs.py as described in (Yi et al. 2010). As PBS requires branch lengths, Fst values were converted using the following equation:

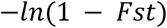

Variants within enhancers with variation between solitary and social populations of *L. albipes* were retained for downstream analysis. Values above the 95^th^ percentile for each group were considered outliers.

## Author Contributions

BMJ and SDK designed and conceptualized the study. Samples were collected by SDK. BMJ, SS, and SMG generated data and data curation was handled by BMJ, AEW, RMS, and MB. SMG, SS, RMS, MB and AEW assembled and annotated the genome with supervision from JE. BMJ and AEW analyzed genomic data. BMJ generated figures and wrote the initial draft of the manuscript. SDK supervised the work. All authors reviewed and edited the final manuscript.

## Acknowledgments

The authors would like to acknowledge T. Simmonds, R. Corpuz, and J. Schrader for library preparation and sequencing support. *Lasioglossum albipes* was sequenced as part of the I5K and USDA-ARS Beenome100 initiative. The US Department of Agriculture, Agricultural Research Service is an equal opportunity/affirmative action employer and all agency services are available without discrimination. This research used resources provided by the SCINet project of the USDA Agricultural Research Service, ARS project number 0500-00093-001-00-D, the Tropical Pest Genetics and Molecular Biology Research Unit in-house appropriated research project number 2040-22430-028-000-D, and the Genetics and Bioinformatics Research Unit in-house appropriated research project number 6066-21310-006-000-D. We also thank the Drosophila Genomics Resource Center (NIH Grant 2P40OD010949) for maintenance of and access to cell lines and the Princeton University Genomic Core staff for sequencing support. Members of the Kocher Lab provided valuable discussion and feedback. This work was supported by Grant DEB 1754476 from the National Science Foundation, and SDK was supported by an NIH New Innovator Award (DP2 2OD027430), the Packard Foundation, and the Pew Biomedical Scholars program. SDK is an HHMI Freeman Hrabowski Scholar.

