## Supplemental Figures for "Repeated shifts in sociality are associated with fine-tuning of highly conserved and lineage-specific enhancers in a socially flexible bee"

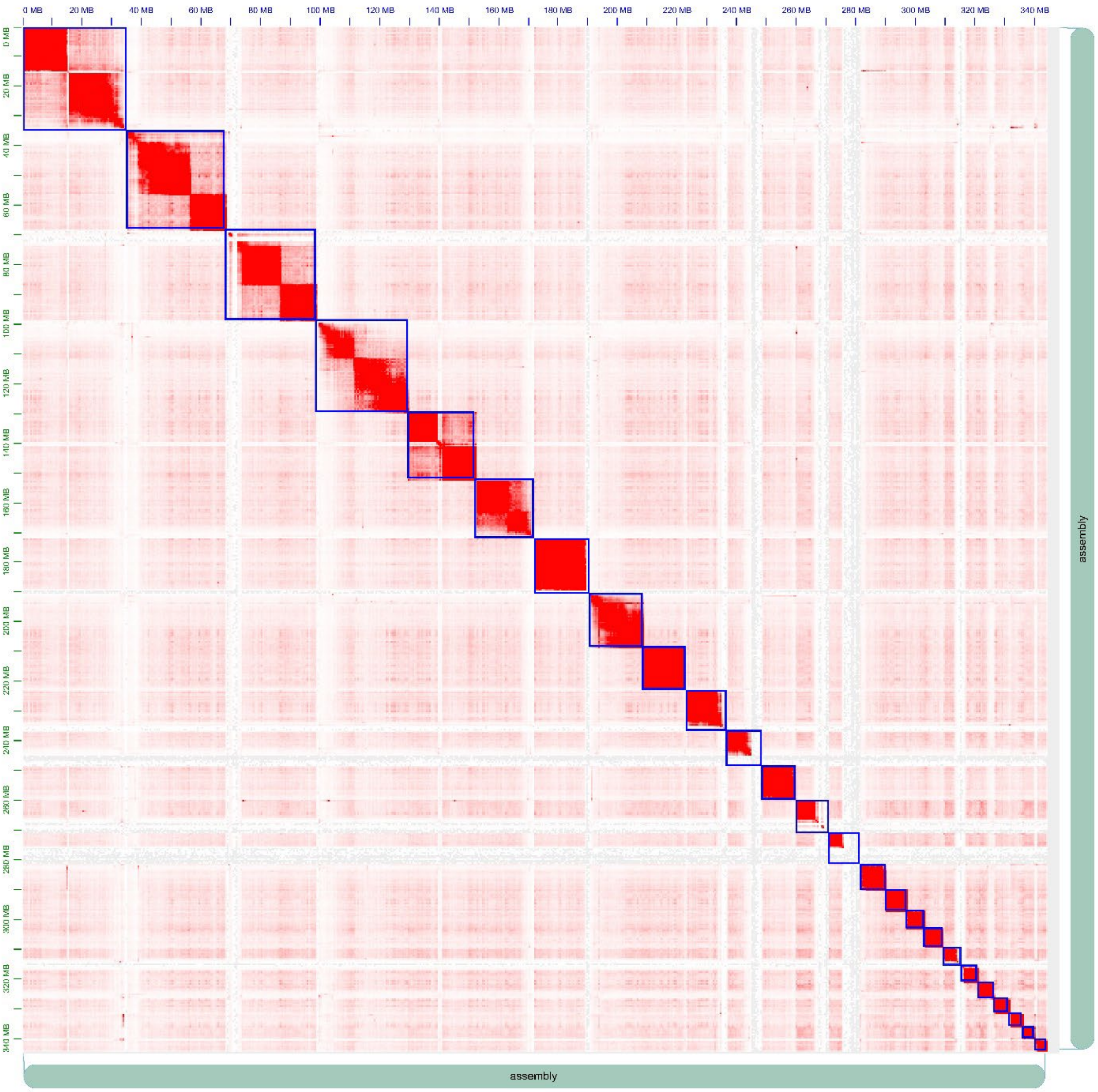


Fig. S1

Updated genome assembly includes contigs placed into 25 chromosomes. Enriched chromosome conformation (HiC) contact map for L¡.albipes assembly. Short 2x150 bp reads from a HiC library generated from one male L¡.albipes was used to assemble the contig assembly into chromosomes based on contact abundance which is visualized as intensity between loci. Blue boxes represent the designated chromosomes.

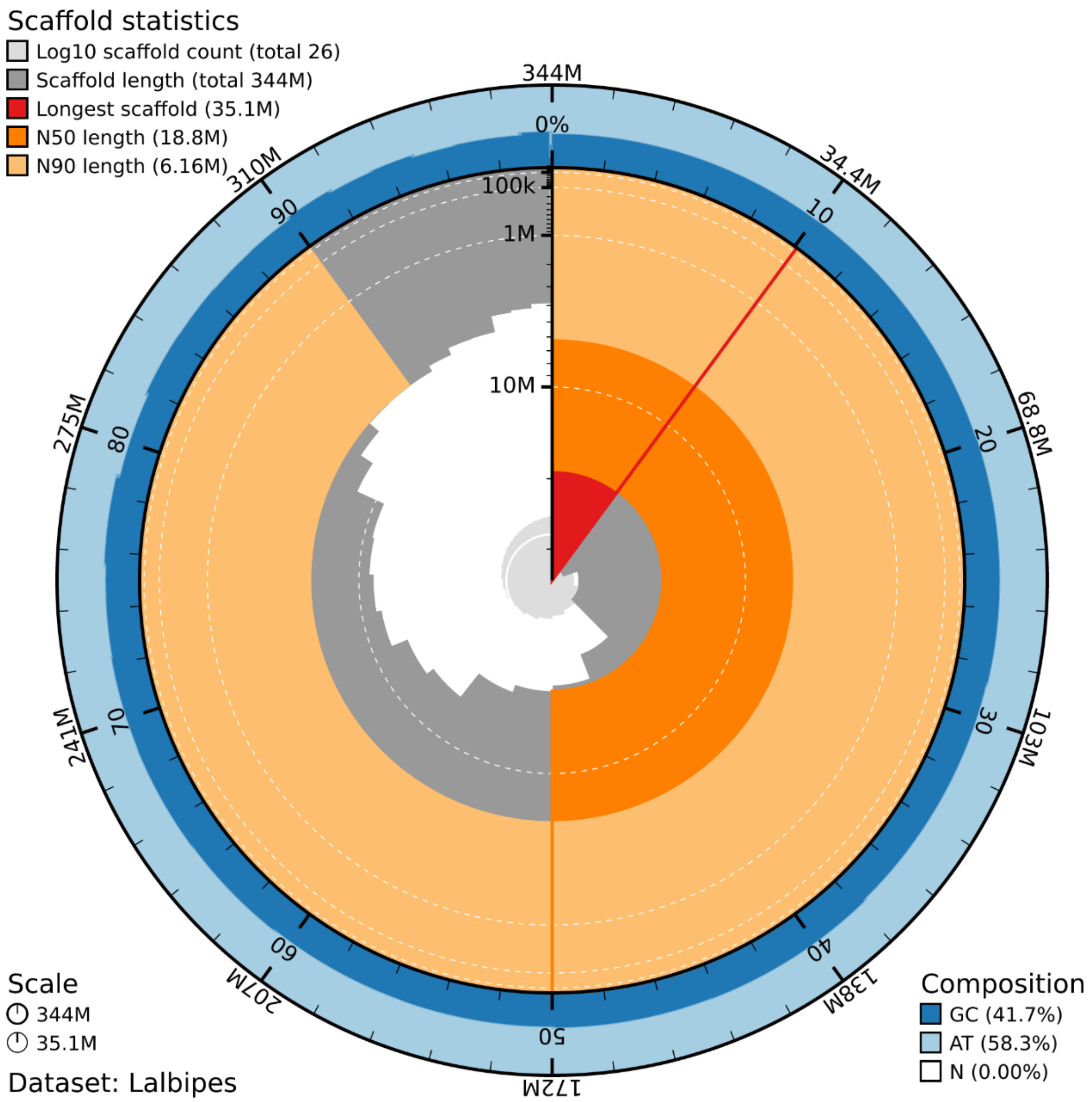


Fig. S2

Assembly metrics for the updated Lasioglossum.albipes.genome. BlobToolKit snail plot of the updated L¡.albipes.assembly after removal of non-arthropod and no-hit contigs. The main plot is divided into bins around the circumference with distribution of sequence lengths shown in dark gray. The plot radius is scaled to the longest sequence present in the assembly (35.052 Mbp, shown in red). Orange and pale-orange arcs show the N50 and N90 sequence lengths (18.823 and 6.162 Mbp), respectively. The pale gray spiral shows the cumulative sequence count on a log scale with white scale lines showing successive orders of magnitude. The blue and pale-blue area around the outside of the plot shows the distribution of GC, AT, and N percentages in the same bins as the inner plot.

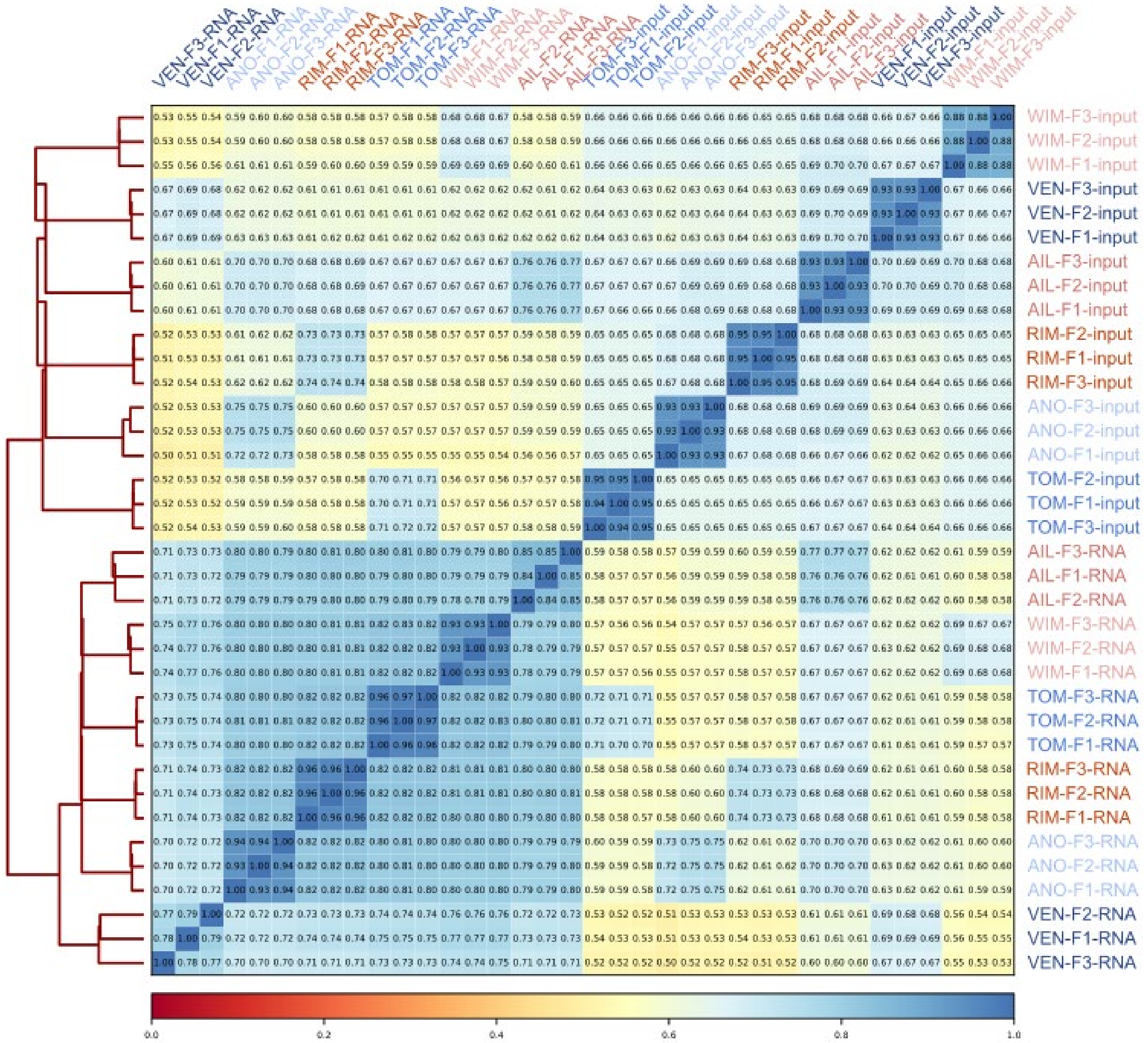


Fig. S3

STARR-seq input and plasmid-derived mRNA libraries are correlated within populations. Heatmap displays Spearman correlation values on read counts for each pairwise comparisons of samples as calculated with deepTools v3.5.5. Dendrogram indicates which samples' read counts are most similar to each other. All mRNA libraries cluster separately from all input samples, and then each biological replicate within each population is most similar to the other replicates for the same population. Input libraries were less correlated (range: 0.6169-0.9501, mean: 0.6916) compared with RNA libraries (range: 0.7022-0.9653, mean: 0.7951).

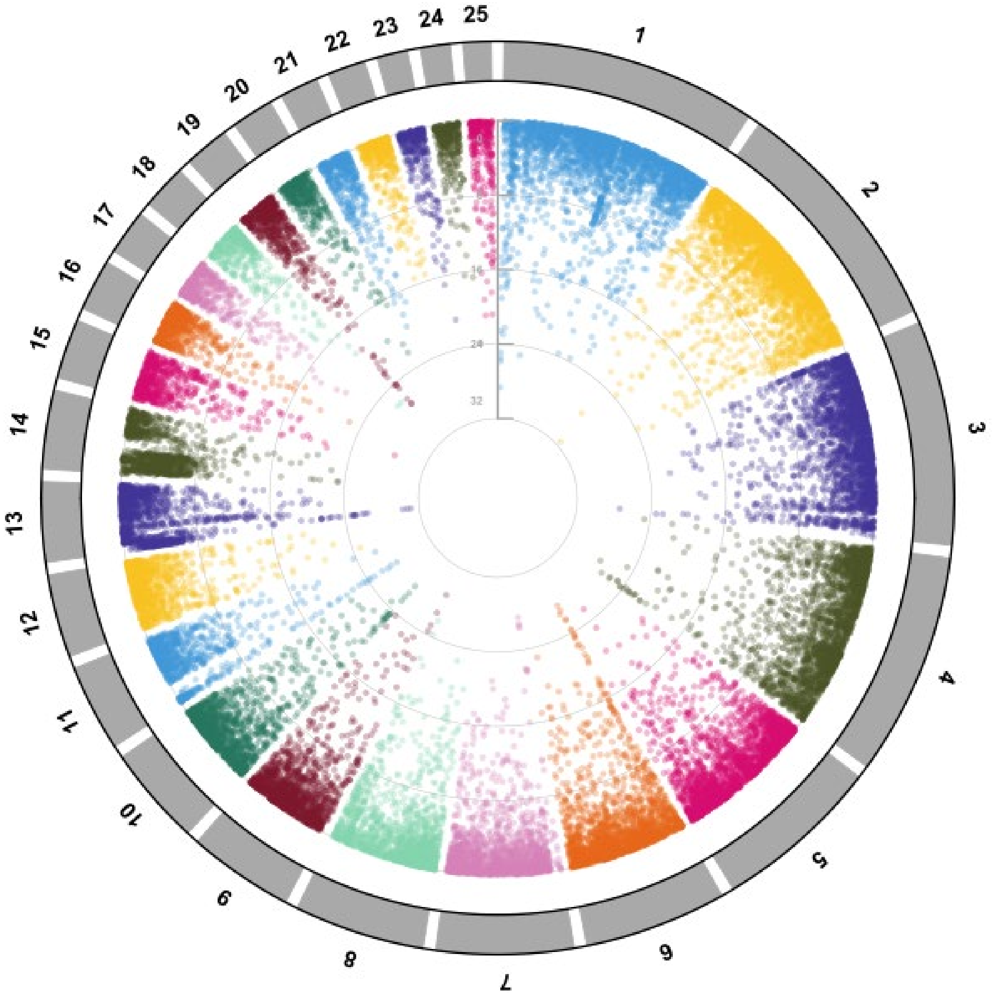


Fig. S4

Identified enhancers were located genome-wide. Circular Manhattan plot displays the log-adjusted p-value of each enhancer identified across populations. Grey bars represent each chromosome, and each filled circle is one enhancer.

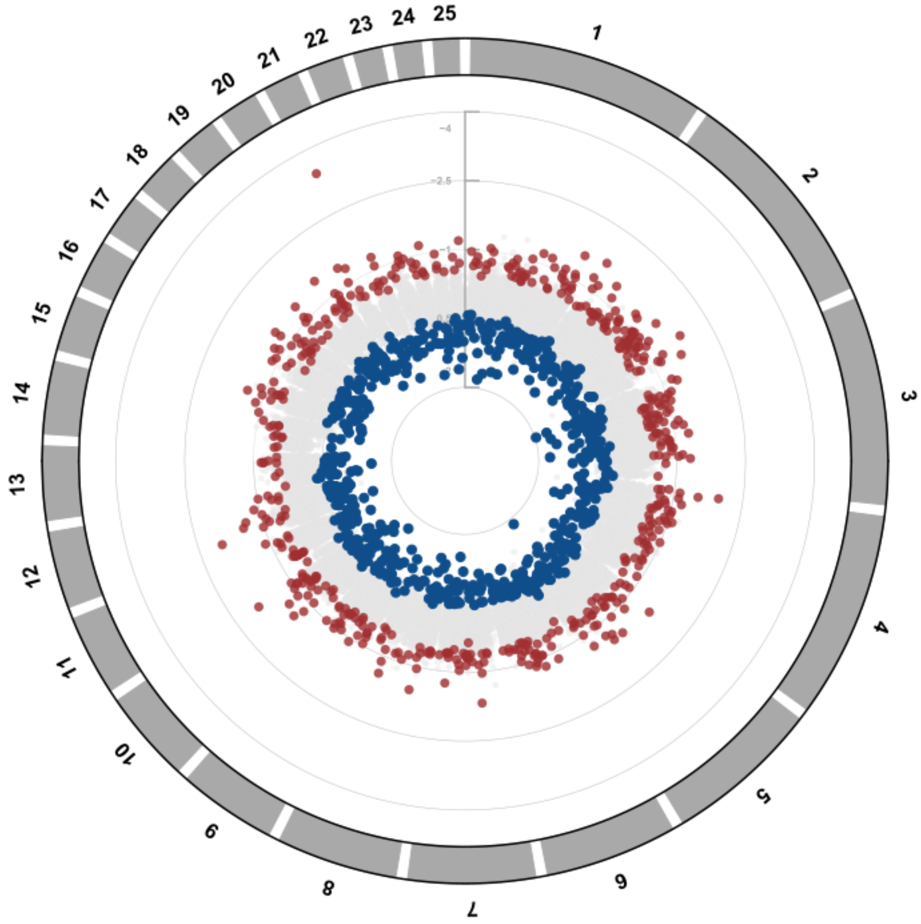


Fig. S5

Enhancers with differential activity between social and solitary populations were identified throughout the genome. Circular Manhattan plot displays the fold-change of STARR-seq enhancer activity between social (red) and solitary (blue) populations. Grey circles indicate enhancers with no difference between social and solitary groups. Grey bars along circumference represent each chromosome, and each filled circle is one enhancer.

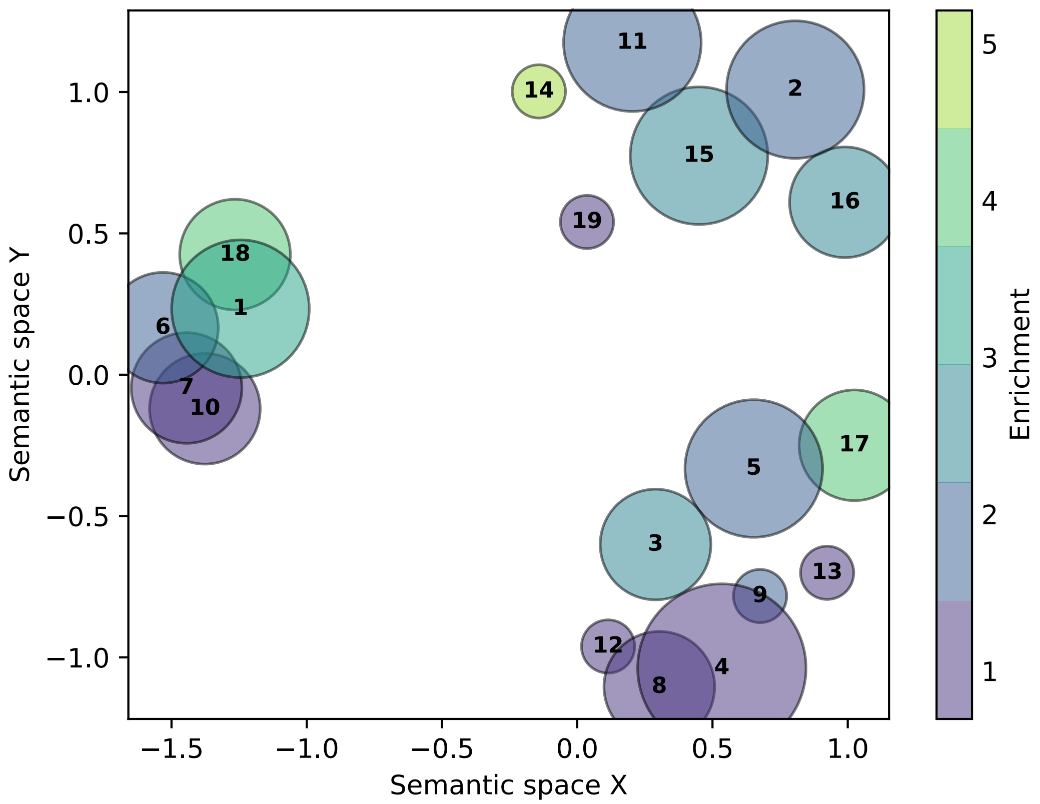


1. eye morphogenesis
2. behavior
3. regulation of protein-containing complex assembly
4. negative regulation of cellular process
5. cell-cell signaling
6. anatomical structure morphogenesis
7. anatomical structure development
8. positive regulation of cellular process
9. regulation of cell population proliferation
10. cellular developmental process
11. axon guidance
12. regulation of cell communication
13. regulation of signaling
14. endothelial cell migration
15. cell-cell adhesion via plasma-membrane adhesion mo...
16. neuron recognition
17. regulation of epithelial cell differentiation
18. establishment of planar polarity
19. nucleic acid metabolic process

Fig. S6

Genes proximal to DAEs are enriched for GO terms related to cellular signaling, morphogenesis, and gene regulation. Input genes included all genes within 10kb of all DAEs and background set was all genes within 10kb of any enhancer. GO enrichment was determined using GOATOOLS (Klopfenstein et al., 2018) and all significant (Benjamini-Hochberg corrected $p<0.05$) terms were clustered by semantic similarity using GO-Figure! (Reijnders & Waterhouse, 2021). Size of each circle is proportional to the specificity of the GO term (more specific = larger), and color indicates enrichment of each GO term in the input set relative to the proportion of genes in the background set.

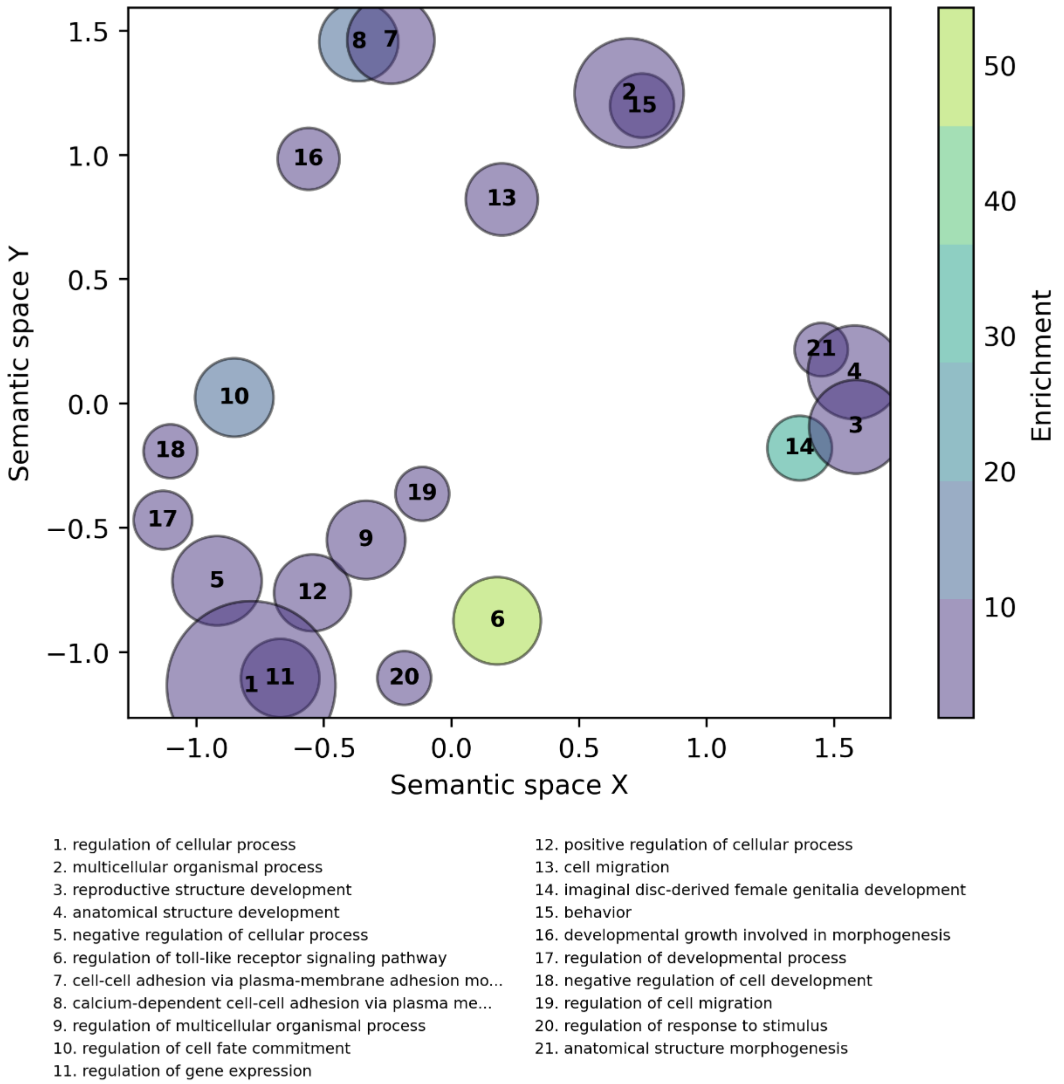


Fig. S7

Genes proximal to conserved DAEs are enriched for GO terms related to cellular and organismal regulation, reproductive and anatomical structure development, multicellular processes, and cell-cell adhesion. Input genes included all genes within 10kb of DAEs active in all 6 populations and background set was all genes within 10kb of any enhancer. GO enrichment was determined using GOATOOLS (Klopfenstein et al., 2018) and all significant (Benjamini-Hochberg corrected $p<0.05$) terms were clustered by semantic similarity using GO-Figure! (Reijnders & Waterhouse, 2021). Size of each circle is proportional to the significant of the GO term (lower adjusted p-value = larger), and color indicates enrichment of each GO term in the input set relative to the proportion of genes in the background set.

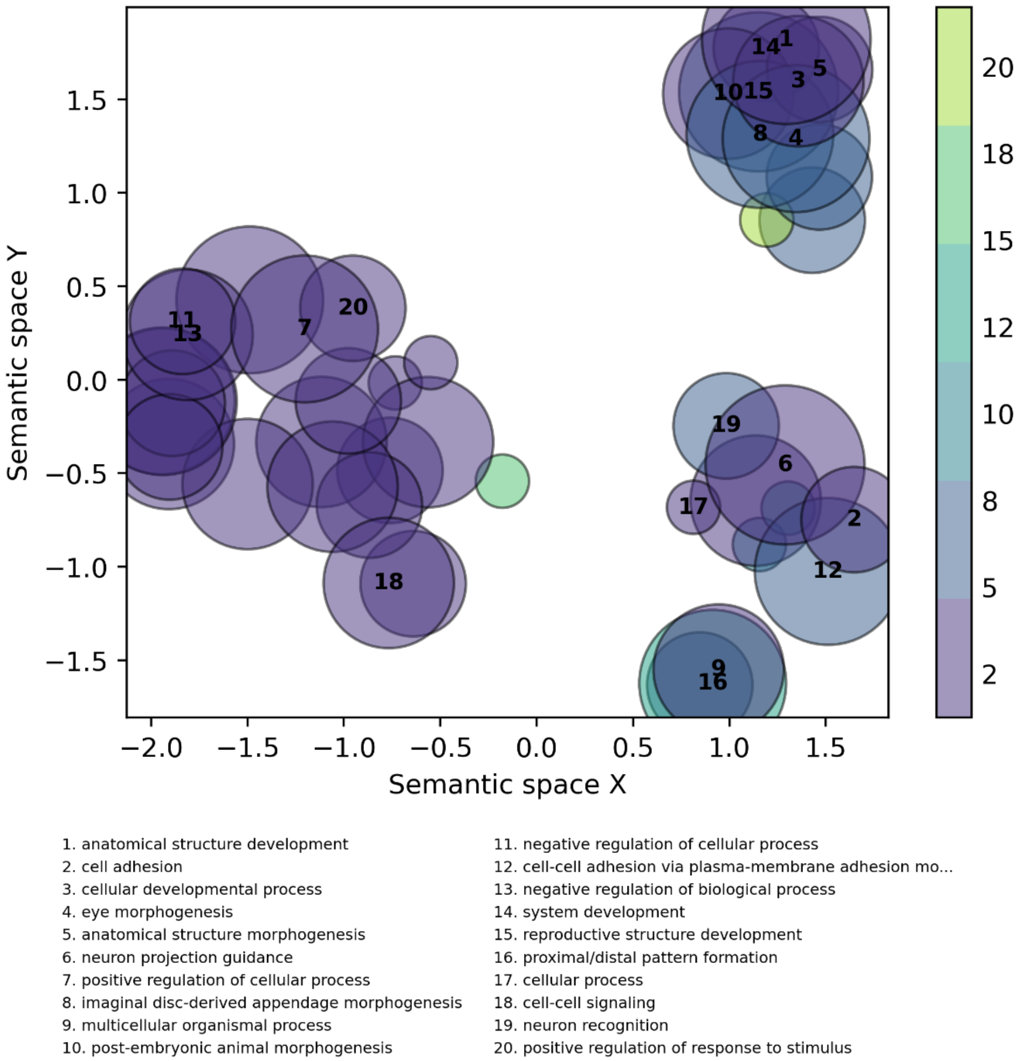


Fig. S8

Genes proximal to both DAEs and fast-evolving CNEEs are enriched for 170 GO terms which cluster semantically into 3 large groups. Input genes included 341 genes that were within 10kb of DAEs and within 10kb of fast-evolving CNEEs between social and solitary lineages of halictid bees (Jones et al., 2023) and background set was all genes within 10kb of any enhancer. GO enrichment was determined using GOATOOLS (Klopfenstein et al., 2018) and all significant (Benjamini-Hochberg corrected $p<0.05$) terms were clustered by semantic similarity using GO-Figure! (Reijnders & Waterhouse, 2021). Size of each circle is proportional to the specificity of the GO term (more specific = larger), and color indicates enrichment of each GO term in the input set relative to the proportion of genes in the background set.

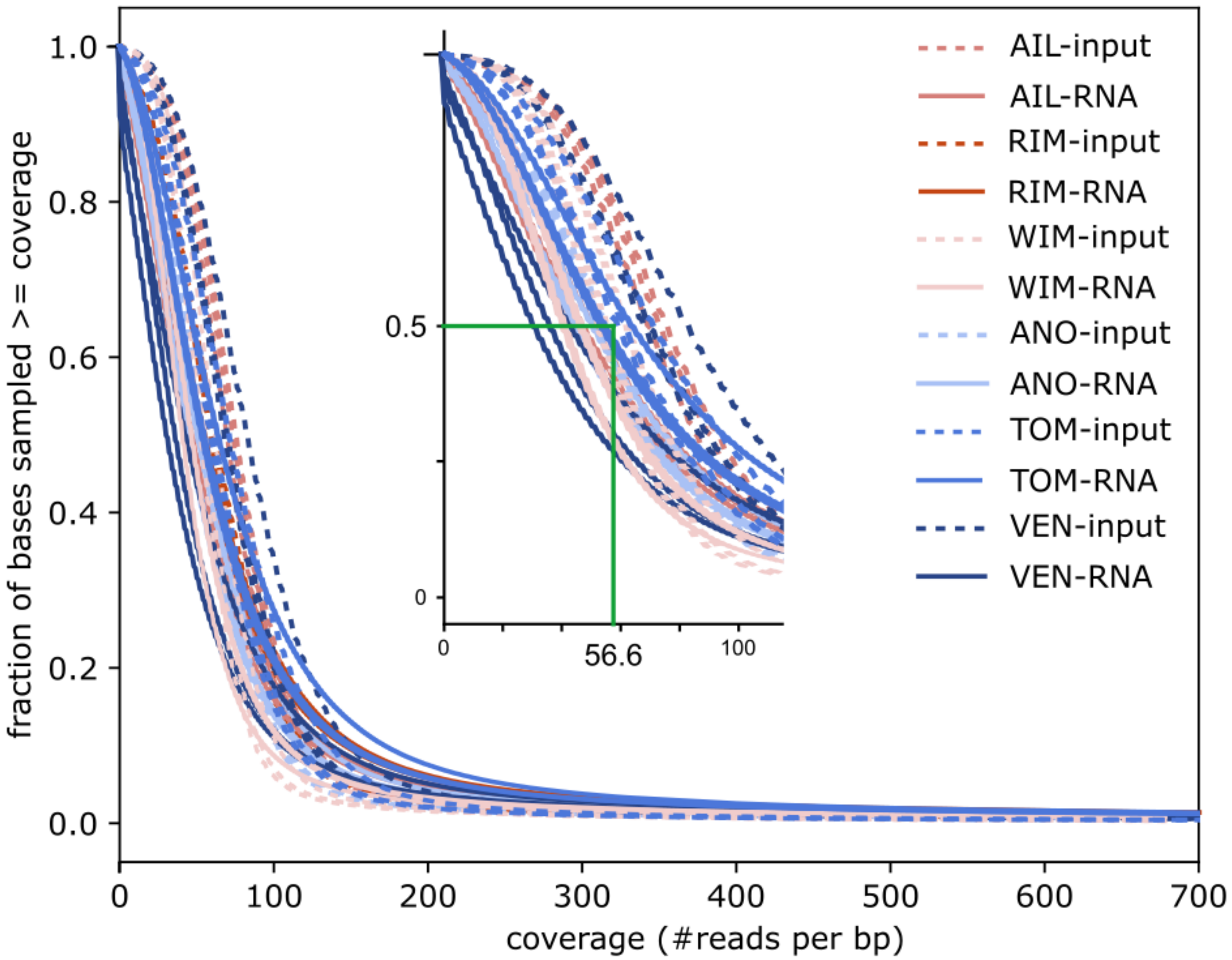


Fig. S9

Genome coverage was similarly high across populations for both STARR-input and STARR-RNA libraries. Reverse cumulative distribution representing the fraction of the genome (y-axis) with a given depth of coverage (x-axis). Inset highlights a fraction of 50%, demonstrating that on average across samples, half of the genome was covered by at least 56.6 reads. 93.3% of the genome was covered by at least 20 reads and 97.5% of the genome was covered by at least 10 reads. Plot was produced with the plotCoverage function of deepTools.
